# Targeting BMI-1 to deplete antibody-secreting cells in autoimmunity

**DOI:** 10.1101/2023.07.27.550912

**Authors:** Jack Polmear, Lauren Hailes, Moshe Olshansky, Maureen Rischmueller, Elan L’Estrange-Stranieri, Anne L Fletcher, Margaret L Hibbs, Vanessa L Bryant, Kim L Good-Jacobson

## Abstract

**Objectives:** B cells drive the production of autoreactive antibody-secreting cells (ASCs) in autoimmune diseases such as Systemic Lupus Erythematosus (SLE) and Sjögren’s syndrome, causing long-term organ damage. Current treatments for antibody-mediated autoimmune diseases target B cells or broadly suppress the immune system. However, pre-existing long-lived ASCs are often refractory to treatment, leaving a reservoir of autoreactive cells that continue to produce antibody. Therefore, the development of novel treatment methods targeting ASCs is vital to improve patient outcomes. Our objective was to test whether targeting the epigenetic regulator BMI-1 could deplete ASCs in autoimmune conditions *in vivo* and *in vitro*.

**Methods:** Use of a BMI-1 inhibitor in both mouse and human autoimmune settings was investigated. *Lyn^-/-^* mice, a model of SLE, were treated with the BMI-1 small molecule inhibitor PTC-028, before assessment of ASCs, serum antibody and immune complexes. To examine human ASC survival, a novel human fibroblast-based assay was established, and the impact of PTC-028 on ASCs derived from Sjögren’s syndrome patients evaluated.

**Results:** BMI-1 inhibition significantly decreased splenic and bone marrow ASCs in *Lyn^-/-^*mice. The decline in ASCs was linked to aberrant cell cycle gene expression and led to a significant decrease in serum IgG3, immune complexes and anti-DNA IgG. PTC-028 was also efficacious in reducing *ex vivo* plasma cell survival from both Sjögren’s syndrome patients and age-matched healthy donors.

**Conclusion:** These data provide evidence that inhibiting BMI-1 can deplete ASC in a variety of contexts and thus BMI-1 is a viable therapeutic target for antibody-mediated autoimmune diseases.

## INTRODUCTION

Antibody-mediated autoimmune diseases encompass a variety of heterogeneous conditions including Systemic Lupus Erythematosus (SLE) and Sjögren’s syndrome. SLE is the most common autoantibody-mediated disease with a prevalence in Australia of 45.3 per 100,000 people, with women and First Nations people being 5 and 2 times more likely to be diagnosed with SLE, respectively (1). SLE patients suffer from chronic and systemic inflammation driven by deposition of antigen-antibody complexes. Glomerulonephritis, one severe outcome of the hyper-inflammatory state in SLE, results in kidney damage often to the point where dialysis or renal transplant is required (2). Sjögren’s syndrome has a global prevalence between 0.22% and 2.7%, with women also being between 9 to 21 times more likely to be diagnosed (3). Pathology is caused by inappropriate immune activation targeting the salivary and lacrimal glands leading to cell destruction (4). Autoantibodies drive innate immune cell activation, stimulating an inflammatory response (5). Limited ability to secrete saliva and tears causes severe ocular dryness and accelerated dental decay, alongside other more severe antibody-driven symptoms such as cryoglobulinemia and vasculitis (3). While tissue pathology may differ between SLE and Sjögren’s Syndrome, symptoms overlap considerably. Autoantibodies produced by antibody-secreting cells (ASCs) drive pathology in both conditions and therefore are a common therapeutic target.

In the absence of available cures, current treatments include broad-spectrum immunosuppressants, or monoclonal antibody therapies such as rituximab and belimumab which target CD20, a B cell surface marker, and BAFF, a B cell survival factor, respectively (6, 7). These latter treatments deplete B cells, and while limiting the ability to generate more pathogenic plasma cells, do not target pre-existing long-lived ASCs that are a source of autoantibodies (8, 9). Bortezomib, a proteasome inhibitor, can reduce ASC numbers, however, approximately half of all patients receiving Bortezomib cease treatment due to adverse events (10). The elimination of pathogenic autoantibodies is key to reducing symptoms and organ damage, and thus improving quality of life. Therefore, novel therapeutic targets that can directly deplete long-lived ASCs are vital for treatment of antibody-mediated autoimmunity.

Histone modifiers have recently emerged as a promising class of therapeutic agents to modulate the humoral immune system (11–15). Histone modifiers add post-translational modifications to histones to regulate chromatin compaction and consequently gene expression (16). As histone modifiers can drive oncogenesis, this has led to the development of small molecule inhibitors as treatment methods (17, 18).

The polycomb repressive complexes (PRCs) are histone-modifying complexes that are essential for mediating activated B cell and ASC fate (12–14, 19, 20). In a previous study, we demonstrated that BMI-1, a component of PRC1, is involved in murine ASC differentiation and survival post-immunisation or post-infection (13). Furthermore, BMI-1 inhibition restored appropriate ASC development during chronic viral infection (13). *BMI1* gene expression is specifically upregulated in human ASCs, compared to other mature B cell populations (21), and thus it may be a viable molecular target to deplete plasma cells in autoimmune settings. Therefore, we set out to test whether BMI-1 inhibition could limit ASC numbers in antibody-mediated autoimmunity.

## RESULTS

### BMI-1 inhibition reduced ASCs in *Lyn^-/-^* mice

To assess the impact of BMI-1 inhibition on ASCs in an autoimmune environment, *Lyn^-/-^* mice were utilised. *Lyn^-/-^* mice are a model of antibody-mediated autoimmunity that have increased numbers of ASCs, particularly splenic plasmablasts, and elevated autoreactive antibodies (22, 23). The onset of SLE-like disease in *Lyn^-/-^* mice is variable, however, at 6 months of age, nearly all *Lyn^-/-^* mice show visible signs of autoimmunity, have high titres of anti-nuclear antibodies, and have kidney disease (24). Thus, our current study utilised *Lyn^-/-^* mice that were 6 months of age or older to mimic severe disease progression.

To investigate whether targeting BMI-1 could reduce autoreactive ASCs, we used PTC-028, a commercially available BMI-1 inhibitor (25). PTC-028 induces BMI-1 hyper-phosphorylation followed by a reduction in BMI-1 (25). *Lyn^-/-^* mice older than 6 months of age were treated daily with PTC-028 for 12 days, followed by assessment four days later at day 15 (d15); Figure 1A). Splenic ASCs, defined as CD138^high^B220^low^ lymphocytes to encompass both plasmablast and plasma cell subsets, were measured via flow cytometry (Figure 1B). The frequency of ASCs as a percentage of lymphocytes in *Lyn^-/-^* spleen was significantly reduced 2-fold following PTC-028 treatment (Figure 1C). Furthermore, a decrease in the total number of splenic ASCs was also observed, with a PTC-028-dependent 2-fold reduction on d15 (Figure 1D). This ASC decrease was not due to an overall decrease in B cells, lymphocytes or splenocytes, as these populations remained unaffected by PTC-028 treatment (Supplementary Figure 1A-D). Therefore, PTC-028 treatment specifically decreased splenic ASCs in *Lyn^-/-^* mice.

**Figure 1:**
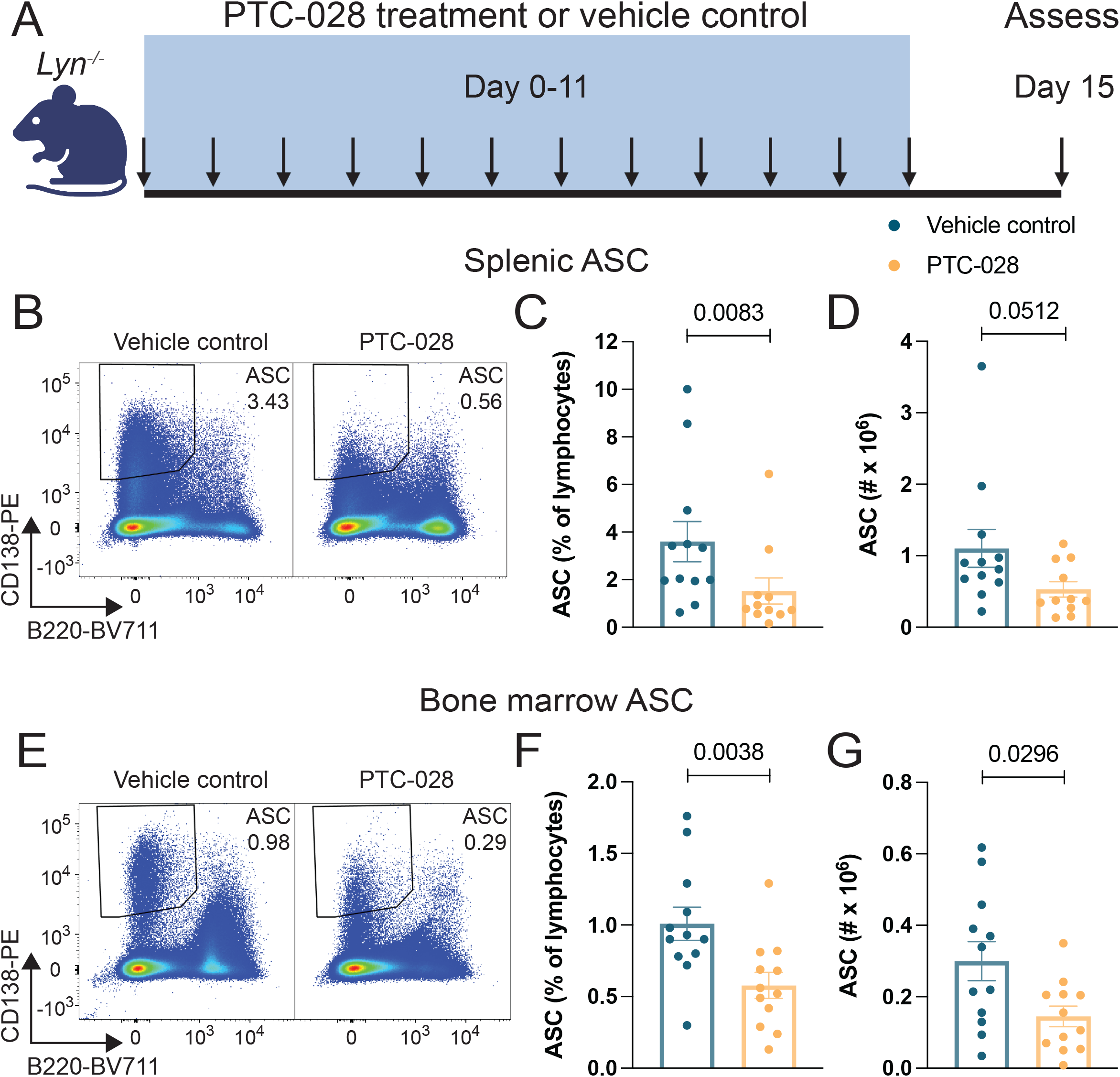
Depletion of ASCs in autoimmune-prone *Lyn^-/-^* mice treated with the BMI-1 small molecule inhibitor PTC-028. A) Schematic of experimental design. *Lyn^-/-^* mice (≥ 6 months old) were treated with PTC-028 or vehicle control once daily on d0 - 11 and then assessed on d15. B) Representative flow cytometric plots of splenic ASCs from *Lyn^-/-^* mice treated with vehicle control or PTC-028. C) Frequency of splenic ASC as a proportion of total lymphocytes and D) total number of splenic ASCs in *Lyn^-/-^* mice treated with vehicle control or PTC-028 on d15. E) Representative flow cytometric plots of bone marrow ASCs from *Lyn^-/-^* mice treated with vehicle control or PTC-028. F) Frequency of bone marrow ASC as a proportion of total lymphocytes and G) total number of bone marrow ASCs in *Lyn^-/-^*mice treated with vehicle control or PTC-028 on d15. Data is presented as the mean +/- SEM with individual data points representing one mouse. Data was combined from three independent experiments. n = 11–12 per group. Mann-Whitney nonparametric two tailed tests were used for statistical analysis.

Bone marrow is enriched with long-lived plasma cells and can host autoreactive ASCs following B cell depletion therapy (26), and therefore bone marrow ASCs were assessed using flow cytometry (Figure 1E). Similar to splenic ASCs, PTC-028 treatment caused a 2-fold reduction at d15 in both frequency and total number of *Lyn^-/-^* ASCs per leg (Figure 1F & Figure 1G), while having no significant impact on B cells, lymphocytes or total cellularity within the bone marrow (Supplementary Figure 1E-H). Together, these data reveal that both splenic and bone marrow ASCs were sensitive to PTC-028 treatment in *Lyn^-/-^* mice.

### BMI-1 inhibition reduced serum IgG3 and anti-DNA IgG

While ASCs are the producers of immunoglobulin, antibody itself is the driver of pathology in antibody-mediated autoimmunity. Antibodies cause damage via immune complex-driven inflammation in SLE or activation of innate immune cells in Sjögren’s syndrome (2, 5) and therefore both serum antibodies and immune complexes were assessed. To determine whether PTC-028 led to a reduction in antibody, enzyme-linked immunosorbent assays (ELISAs) were performed on d15 serum samples from *Lyn^-/-^* mice for each immunoglobulin isotype. Total IgG3 was reduced significantly following BMI-1 inhibition (Figure 2A), whereas IgA, IgM, and the other IgG subtypes were unaffected by PTC-028 treatment at d15 (Supplementary Figure 2A-E). Data from age-matched wild-type mice greater than 6 months old similarly showed a large decrease in IgG3 serum levels (Supplementary Figure 2F), suggesting that BMI-1 inhibition may specifically target IgG3-secreting ASCs in aged mice. In addition, circulating IgG3 immune complexes were significantly decreased in *Lyn^-/-^* mice treated with PTC-028 compared to vehicle control mice (Figure 2B).

**Figure 2:**
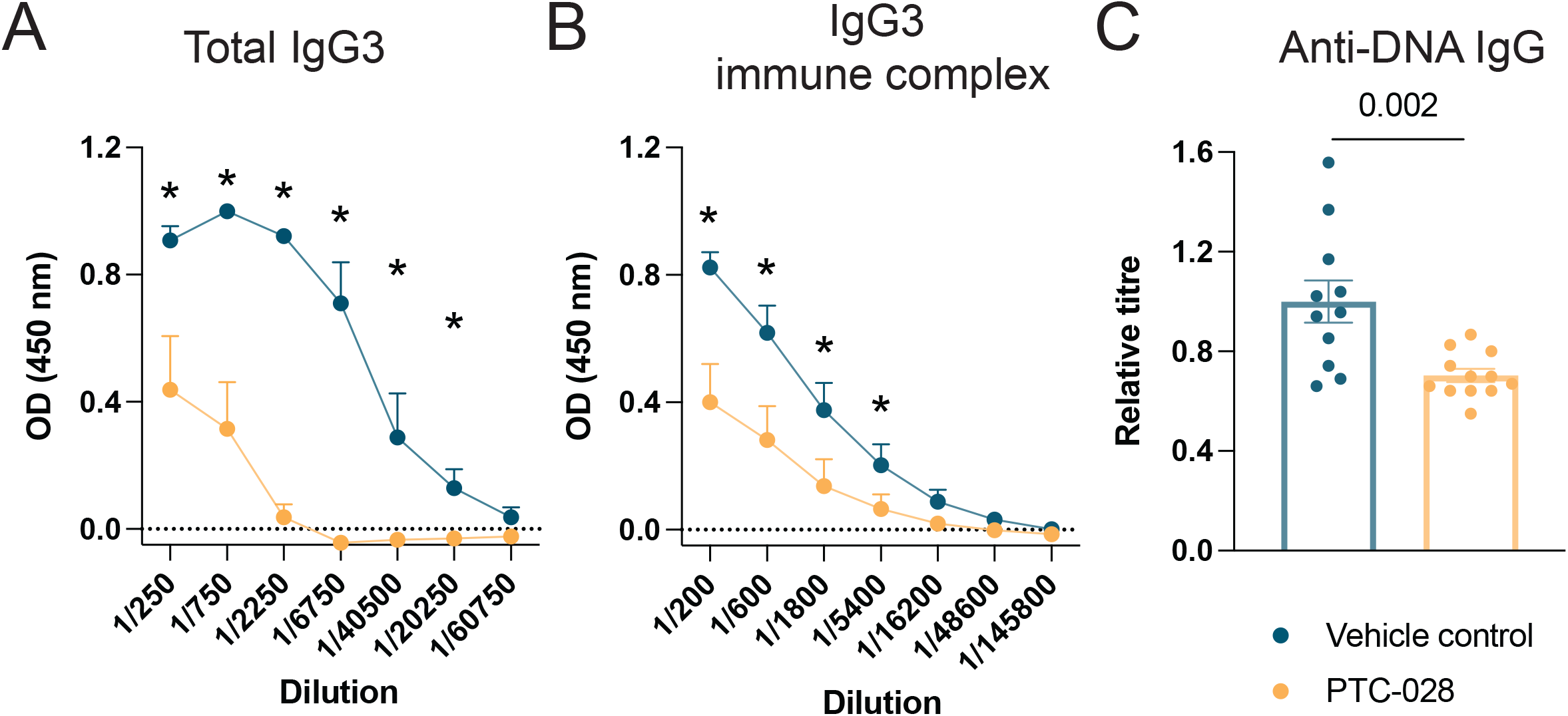
BMI-1 inhibition reduced total IgG3 and anti-DNA IgG in *Lyn^-/-^* mice. A) Assessment of serum antibody by ELISA. Total IgG3 in *Lyn^-/-^* mice treated with vehicle control or PTC-028 once daily on d0 - 11 and then assessed on d15. Data shown is representative of three independent experiments. n=4 per group. B) IgG3 immune complexes and C) anti-DNA IgG in *Lyn^-/-^* mice treated with vehicle control or PTC-028 once daily on d0 - 11 and then assessed on d15. n=11–12 per group. Data was combined from three independent experiments. Mann-Whitney nonparametric two tailed tests was used for statistical analysis. *P≤0.05.

DNA is a major autoantigen in many systemic antibody-mediated autoimmune diseases, particularly in SLE (2). *Lyn^-/-^* mice have high amounts of pathogenic anti-DNA IgG2b, IgG2c and IgG3 (27). PTC-028 treatment in *Lyn^-/-^* mice significantly reduced anti-DNA IgG 1.4-fold compared to vehicle control (Figure 2C). Together, these results demonstrated that BMI-1 inhibition reduced IgG3 immune complexes and pathogenic anti-DNA antibodies in *Lyn^-/-^* mice. Thus, BMI-1 inhibition may be a promising therapeutic strategy to reduce autoreactive ASCs *in vivo* in antibody-mediated autoimmune disease.

### BMI-1 inhibition regulated cell cycling genes in ASCs

RNA sequencing was performed to elucidate the molecular changes induced by PTC-028 in ASCs in *Lyn*^-/-^ mice. ASCs were sort-purified from *Lyn^-/-^* mice on d15 following in vivo PTC-028 or vehicle control treatment. RNA sequencing analysis revealed a total of 587 differentially expressed genes (DEGs) between the PTC-treated and vehicle-treated ASCs. While 540 genes were upregulated in ASCs treated with PTC-028, only 47 genes were downregulated (Figure 3A).

**Figure 3:**
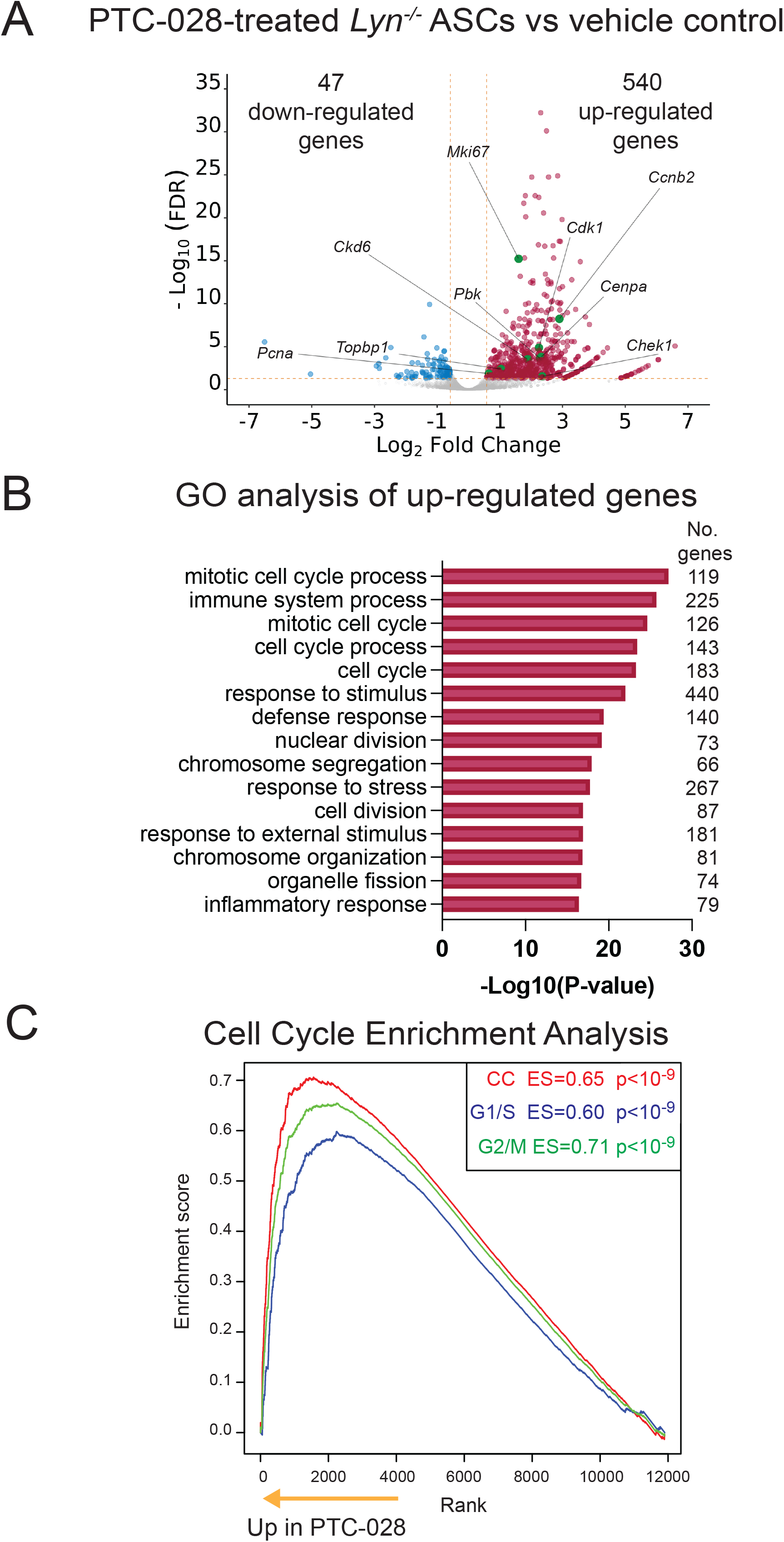
BMI-1 inhibition regulated cell cycle genes in ASCs isolated from *Lyn^-/-^* mice. RNA sequencing was performed on splenic ASCs (B220^low^CD138^hi^) sort-purified from *Lyn^-/-^* mice on d15 following treatment with PTC-028 or vehicle control from d0-11. A) Volcano plot comparing average gene expression in ASCs from *Lyn^-/-^* mice treated with either PTC-028 or vehicle control. False discovery fate (FDR) (minimum cut-off = 0.05) is compared to fold change (Minimum cut-off = +/- 1.5). Upregulated genes are displayed in red, downregulated genes are displayed in blue and grey represents non-significant results. Labelled genes (in green) are genes associated with the cell cycle that are regulated by BMI-1. B) Gene ontology (GO) analysis on upregulated genes showing the top 15 most statistically enriched biological processes. The number of enriched genes (no. genes) is listed for each process. C) Cell cycle enrichment analysis showing enrichment scores in our RNA sequencing data set for genes associated with the cell cycle (CC), G1/S phase and G2/M phase. n=2 per group.

Gene ontology (GO) analysis was performed to identify affected biological processes based on the gene clusters that were enriched in our dataset. As there were few downregulated genes, we focused our analysis on the upregulated gene set. 119 of the upregulated genes in our drug-treated group were found to be involved in the mitotic cell cycle process (p=6.7×10^-28^) (Figure 3B). Furthermore, 6 out of the top 15 predicted biological processes were related to cell cycling. Gene Set Enrichment Analysis (GSEA) also revealed an enrichment for genes involved in cell cycling (Enrichment score (ES)=0.65, p<10^-9^) (Figure 3C), including genes associated with G1/S phase (ES=0.60, p<10^-9^) as well as G2/M phase (ES=0.71, p<10^-9^). This result was consistent with previous studies that have identified BMI-1 as a key regulator of the cell cycle in several other disease and cellular contexts (28–32). We therefore interrogated our dataset for genes which are known to be directly or indirectly regulated through BMI-1 and are involved in cell cycling. Indeed, key regulators of cell cycle processes which have been identified as BMI-1 targets, such as *Cdk1*, *Cdk6*, *Chek1*, *Ccnb2*, *Topbp1*, *Pbk*, *Cenpa*, *Mki67*, *Pcna*, were upregulated in *Lyn^-/-^* ASCs following PTC-028 treatment (Figure 3A). Thus, RNA-sequencing analysis revealed BMI-1 inhibition caused altered gene expression of cell cycle regulators in ASCs from *Lyn^-/-^* mice.

### BMI-1 inhibition limited the survival of ASCs derived from Sjögren’s syndrome patients

For BMI-1 inhibition to be a viable treatment strategy for antibody-mediated autoimmunity it must be able to deplete human ASCs in autoimmune contexts. Therefore, we established an *ex vivo* assay to assess the survival of human ASCs sort-purified from human peripheral blood mononuclear cells (PBMC) obtained from Sjögren’s syndrome patients. ASCs are increased in the peripheral blood of Sjögren’s syndrome compared to healthy controls (33, 34) and this is correlated with disease severity, potentially indicating a higher prevalence of autoreactive ASCs (34). The Sjögren’s syndrome patients who participated in this study had a range of comorbidities, serum IgG concentrations, and disease severity (Supplementary Table 1). All patients were positive for antibodies against SSA/Ro, a key autoantigen in Sjögren’s syndrome (4). All patients tested for rheumatoid factor were positive (Supplementary Table 2; one patient was not tested). Of note, two thirds of patients had been or were on immunomodulatory agents.z

We first established and optimised a 6 day-long fibroblast-based survival assay for human plasma cells. Sort-purified ASCs were cultured in the presence of human tonsil fibroblast supernatant supplemented with APRIL and IL-6, and treated with either PTC-028 or a DMSO vehicle control. Following 6 days of culture, ASC viability was assessed by flow cytometry (Figure 4A). Initially, age-matched healthy donor samples (Supplementary Table 2) were used to assess the impact of BMI-1 inhibition on human ASCs. PTC-028 treatment led to a 4-fold reduction in live ASCs, confirming the susceptibility of human ASCs to BMI-1 inhibition (Figure 4B). Next, we assessed the ability of BMI-1 inhibition to reduce ASCs isolated from autoimmune patients. ASCs isolated from Sjögren’s syndrome patients showed a 2-fold decrease in following PTC-028 treatment compared to the vehicle control (Figure 4C). Despite the heterogeneity in disease severity and current treatment regimen in patients, PTC-028 treatment consistently demonstrated a significant reduction in ASC survival across all samples. Together, these data confirm that using a BMI-1 inhibitor can reduce the survival of human ASCs derived from autoimmune patients.

**Figure 4:**
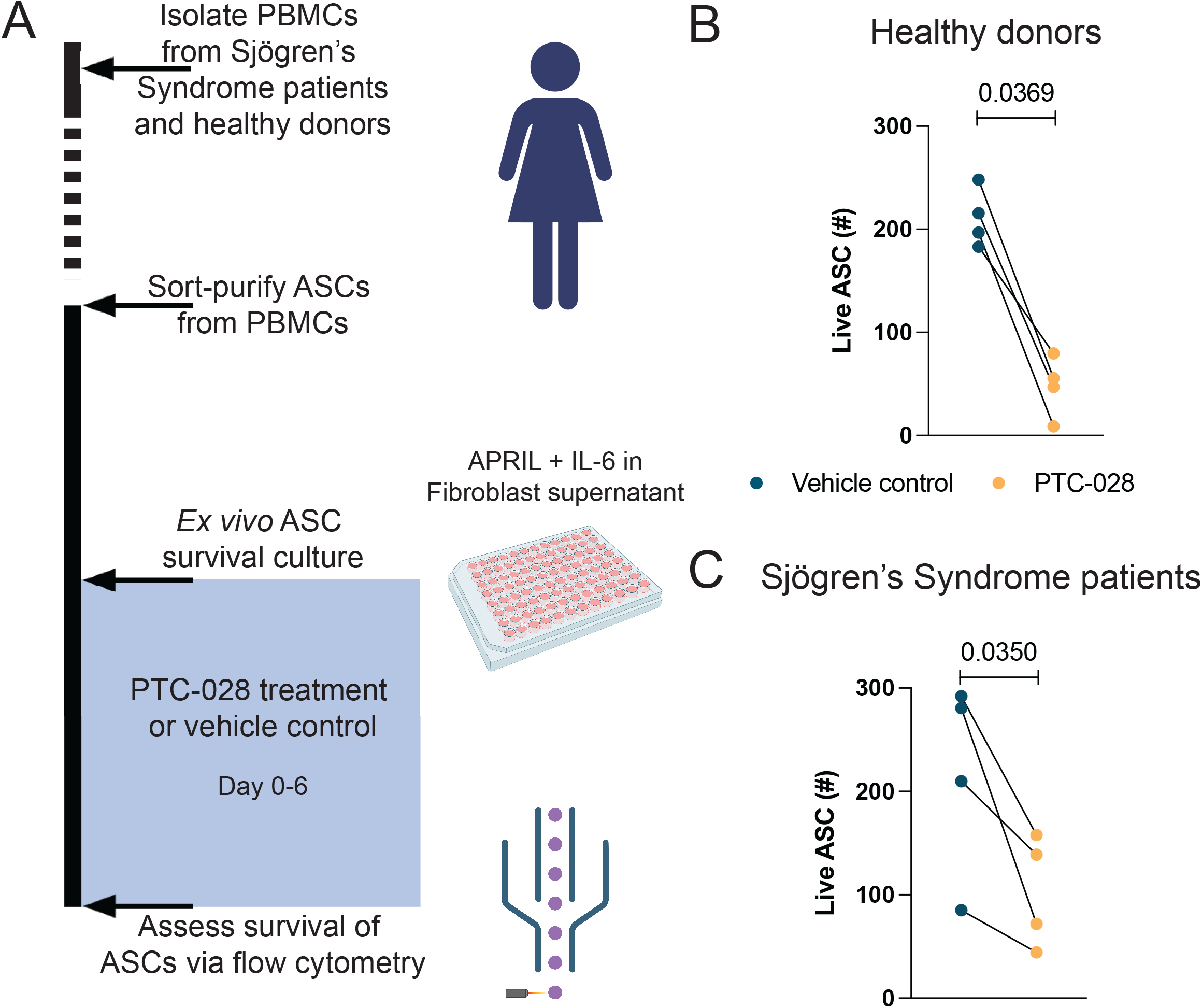
BMI-1 regulated survival of ASCs isolated from Sjögren’s Syndrome patients. A) Schematic of experimental design. ASCs (CD20^-^CD3^-^CD14^-^CD16^-^CD19^+^CD27^hi^CD38^hi^) were sort-purified from *ex vivo* PBMCs isolated from either Sjögren’s Syndrome patients or aged-matched healthy donors. ASCs were cultured in human tonsil fibroblast supernatant with IL-6 and APRIL for 6 days in the presence of PTC-028 or a DMSO vehicle control. Viability of ASCs was analysed using flow cytometry. Total number of live ASCs from B) Healthy donors and C) Sjögren’s Syndrome patients treated with vehicle control or PTC-028. Data is represented as the mean from each biological replicate with connected data points originating from the same donor. n=4 per group. Ratio paired t test was used for statistical analysis.

## DISCUSSION

The current treatment strategy for SLE is based on broad immunosuppression using drugs such as hydroxychloroquine, glucocorticoids and methotrexate (35). When flairs occur or in refractory SLE, drugs specifically targeting B cells are often used (35). Rituximab is typically administered weekly to fortnightly for 2 – 4 weeks, which has shown promising results in decreasing disease activity (36). Treatment for Sjogren’s syndrome is typically based on tear and saliva substitutes in addition to broad immunosuppression in patients with systemic manifestations (37). The use of rituximab in Sjogren’s syndrome initially showed promise, however larger clinical trials have not met their primary outcomes (38). Rituximab does not target existing plasma cells as they only express low levels of surface CD20 (9). Here, we demonstrate that treatment of *Lyn^-/-^* mice with PTC-028, a BMI-1 inhibitor, depleted splenic and bone marrow ASCs and significantly reduced serum IgG3 and anti-DNA IgG. Additionally, BMI-1 inhibition was able to reduce human ASCs obtained from either healthy individuals or Sjögren’s syndrome patients. With clinical trials currently running to assess safety and pharmacokinetics in the context of solid tumour treatment (clinical trial NCT02404480), BMI-1 inhibitors could be a novel treatment to limit the production of autoantibodies in patients with systemic autoimmune disease, particularly those who are not responding well to standard treatments. Therefore, the use of BMI-1 inhibition may improve disease outcomes in patients, particularly those with high levels of long-lived ASCs.

RNA sequencing of ASCs from *Lyn^-/-^* mice revealed 587 DEGs, with the vast majority of genes upregulated post-BMI-1 targeting. This is consistent with the role of BMI-1 in PRC1, a complex which deposits a repressive histone mark H2AK119ub1 (39), and consistent with conditional deletion of BMI-1 in a viral infection model (13). Perturbations in cell cycle genes indicate BMI-1 inhibition alters cell cycle progression. This finding is in line with the role of BMI-1 as a cell cycle regulator. For example, BMI-1 has been shown to alter cell cycle progression via repression of the INK4a locus which inhibits CDK4/6, key regulators of the cell cycle (28, 29). Accordingly, *Cdk6* was upregulated following BMI-1 inhibition in *Lyn^-/-^* mice. Previous work also revealed that BMI-1 overexpression can suppress proliferation and cell cycle regulator genes such as *Mki67*, *Pcna* and *Ccnb2* (30), which were upregulated in our dataset upon BMI-1 inhibition. Our data showed an enrichment of genes associated with both G2/M and G1/S phases of the cell cycle, which in other contexts has led to apoptosis (26) and again highlighted the dysregulation of the cell cycle in PTC-028-treated ASCs. However, commercially available inhibitors such as PTC-028 and PTC-596 can have off target effects in addition to BMI-1-dependent effects on cell biology. Thus, given the importance of targeting BMI-1-expressing cells for therapeutic intervention, there is a need for new drug development to specifically inhibit BMI-1 expression.

IgG2b, IgG2c and IgG3 are the main drivers of pathology in *Lyn^-/-^*mice (27). PTC-028 treatment in *Lyn^-/-^* mice caused a significant decrease in the level of total serum IgG3 and IgG3^+^ immune complexes. While our data demonstrated that ASCs were in part dependent on BMI-1 for survival, which Ig isotype is most affected by the absence or inhibition of BMI-1 appears to be context dependent. *Bmi1* can be detected in IgA, IgM and IgG ASCs (13). Targeting BMI-1 in 3-month-old mice immunised with NP-KLH or infected with LCMV reduced IgG1 and IgG2c, respectively (13). In contrast, BMI-1 inhibition in aged *Lyn^-/-^* mice or aged naïve wild-type mice decreased IgG3. This discrepancy could be due to the increase in IgG3 that occurs with age in C57/BL6 mice (40). Taken together, we propose that the effect of BMI-1 may be most prominent in the primary IgG subclass dominating the specific immune response at the time.

Mouse IgG3 can form larger immune complexes which was demonstrated to be dependent on the IgG3 CH2 domain (41). Mouse and human IgG subclasses are structurally and functionally distinct (42), therefore, identifying the impact of BMI-1 inhibition on human IgG subclasses is an important future direction. IgG3 also has an increased distance between the Fab arms, which may also contribute to increased immune complex formation (43). Furthermore, mouse IgG3 is also a potent activator of complement (41). Immune complexes and complement activation are major drivers of inflammation in many antibody-dependent autoimmune diseases, particularly SLE (2, 4). Therefore, targeting IgG3 may alleviate some inflammation caused by autoreactive antibodies. This is supported by IgG3 causing nephritis in MRL/lpr mice (44). The significant reduction in ASCs and serum IgG3 post-BMI-1 inhibition was concomitant with a corresponding significant decrease in IgG3 immune complexes and anti-DNA IgG, thus underscoring the potential to use BMI-1 small molecule inhibitors to deplete immunopathogenic ASCs and alleviate tissue pathology in autoimmune disease.

In summary, our study demonstrates that BMI-1 inhibition significantly reduces ASCs from autoreactive environments in both mice and humans, establishing a proof of principle that BMI-1 is a viable therapeutic target to deplete ASCs in antibody-dependent autoimmune diseases.

## METHODS

### Mice

*Lyn^-/-^* mice were housed at the Monash Animal Research Platform. All experiments were ethically reviewed and approved by the Monash University Animal Ethics Committee. Experiments were performed in accordance with NHMRC guidelines for the care and use of laboratory animals. Male and female *Lyn^-/-^* mice were aged to at least 6 months prior to treatment. PTC-028 (Sapphire Bioscience) was dissolved in 0.5% Methyl cellulose (Sigma-Aldrich) 1% Tween 80 (MPBio) in MilliQ water. Age-matched and sex-matched *Lyn^-/-^* mice were treated with 15mg/kg of PTC-028 or vehicle control (0.5% Methyl cellulose 1% Tween 80 in MilliQ water) daily for 12 days. Mice were humanely euthanised by hypercapnia. Spleens, hind-leg bones and blood were collected for analysis.

### Human Samples

Peripheral blood mononuclear cells (PBMCs) were isolated from fresh whole blood by density gradient Ficoll-Leucosep centrifugation and cryopreserved in liquid nitrogen. Patients with a diagnosis of Sjogren’s Syndrome were recruited from the Rheumatology Unit, The Queen Elizabeth Hospital, Adelaide. Unrelated healthy donors were recruited from the Volunteer Blood Donor Registry (WEHI). Written, informed consent was obtained from all participants prior to inclusion in the study, and all procedures performed involving human participants were in accordance with the ethical standards of local Human Research Ethics Committees at Central Adelaide Local Health Network, WEHI and Monash and with the 1964 Helsinki declaration and its later amendments or comparable ethical standards. Sjögren’s Syndrome patients were aged between 32 and 67 years old (mean of 57 years old) and included 6 females (Supplementary Table 1). Healthy donors were aged between 33 and 67 years old (mean of 58 years old) and included 6 females (Supplementary Table 2).

### Flow cytometry and Fluorescent-Activated Cell Sorting (FACS)

Single-cell suspensions were prepared and resuspended in PBS 2% FCS. Cell viability was measured using Fixable Viability Stain FVS780 (BD). Mouse cells were incubated with B220 (RA3-6B2, BioLegend) and CD138 (281-2, BioLegend), while human cells were incubated with CD20 (2H7, BD Biosciences), CD19 (SJ25C1, BioLegend), CD27 (M-T271, BioLegend), CD38 (HIT2, BD Biosciences), CD3 (UCHT1, BioLegend), CD14 (HCD14, BioLegend) and CD16 (3G8, BioLegend). Antibody cocktails included normal rat serum (Sigma-Aldrich) and FcγRII/III (2.4G2 supernatant). Live ASCs were sort-purified using an Influx (BD). For flow cytometric analysis, human cells were fixed using Cytofix Fixation Buffer (BD). Multiparameter flow cytometric analysis was performed on a Fortessa X20 flow cytometer (BD) with data acquired using FACSDiva software (BD). Data analysis was performed with FlowJo v10 (TreeStar).

### Isotype specific ELISAs

96-well high-binding ELISA plates (Sarstedt) were coated overnight at 4°C with either goat anti-mouse IgA, goat anti-mouse IgM, goat anti-mouse IgG1, goat anti-mouse IgG2b, goat anti-mouse IgG2c or goat anti-mouse IgG3 (Southern Biotech). Plates were blocked for 1 hr using PBS 1% BSA, washed with PBS 0.05% Tween 20 followed by distilled water, and loaded with serially diluted serum samples for 3 hr at 37°C. Plates were washed prior to adding goat anti-mouse IgA-HRP, goat anti-mouse IgM-HRP, goat anti-mouse IgG1-HRP, goat anti-mouse IgG2b-HRP, goat anti-mouse IgG2c-HRP or goat anti-mouse IgG3-HRP (Southern Biotech) and incubated for 1 hr at 37°C. Plates were washed and then developed using OPD substrate solution (Sigma-Aldrich). Optical density was measured at 450 nm using a FLUOstar OPTIMA (BMG Labtech). Average blank optical density (OD) values were subtracted from sample raw OD values.

### Immune complex assay

Immune complexes from mice sera were precipitated by incubating serum in an equal volume of 5% PEG6000 in PBS overnight at 4°C. This solution was then diluted 1:3 with 2.5% PEG6000 in RPMI and centrifuged at 2000g for 30 minutes at 4°C. IgG3 immune complexes were then measured via ELISA as described.

### Anti-DNA IgG ELISA

96-well Maxisorp Immunoplates (Thermo Scientific Nunc) were coated overnight at 4°C with 50 μg/ml of calf thymus DNA (Sigma Aldrich) diluted in autoclaved PBS 1mM EDTA (50 μl/well). Plates were washed twice with PBS and blocked for 1 hour at room temperature with 100 μl assay diluent (BD). Plates were washed with PBS before adding sera diluted in assay diluent, incubated for 2 hours at room temperature (50 μl/well). A serial dilution of an in-house, high anti-dsDNA IgG reference serum was used to generate a standard curve. Plates were washed six times with PBS 0.05% Tween-20 (Sigma-Aldrich) and incubated for 1 hour at room temperature with horseradish peroxidase–conjugated goat anti–mouse IgG (Southern Biotech) diluted 1:4000 with assay diluent. Plates were washed six times with PBS-tween and developed by adding 50 μl/well of tetramethylbenzidine substrate (BD), with reaction stopped using 1.5M H2SO4. Plates were read at 450 nm, corrected at 570 nm, using the MultiSkan GO microplate spectrophotometer (Thermo Fisher Scientific). Relative titres were determined by plotting the optical density against the standard curve. All samples were normalised to the mean of the vehicle control.

### RNA sequencing and bioinformatics analysis

#### RNA sequencing

ASCs (FVS780^-^B220^low^CD138^hi^) were sort-purified from biological replicates of *Lyn^-/-^* mice treated with either vehicle control or PTC-028. One biological replicate in each group (vehicle or PTC-028) was obtained by pooling sort-purified ASCs from two individual mice due to low cell number. ASCs were lysed using RLP buffer (QIAGEN). RNA was extracted using the RNeasy Plus Micro Kit (QIAGEN) according to the manufacturer’s protocol, removing genomic DNA by using a gDNA eliminator spin column (QIAGEN). Total mRNA quantity and quality was measured using a Qubit Fluorometer (Invitrogen) and Bioanalyzer (Agilent). Total RNA was enriched for mRNA using poly-A enrichment before fragmentation and cDNA synthesis. RNA sequencing was performed by AZENTA Life Sciences on the Illumina Novaseq platform in a 150-base pair paired end configuration.

#### Analysis of RNA sequencing

Paired end reads in fastq format were trimmed of adaptors with TrimGalore version 0.6.10 (cutadapt version 4.3). Meanwhile read quality was assessed with FastQC. Adaptor-trimmed reads were aligned to the mm39 genome (downloaded from the UCSC Genomic Browser) with hisat2 version 2.2.1. Duplicated alignment pairs were marked and removed with MarkDuplicates from Picard Tools version 2.18.0, and alignments with mapping quality < 3 were removed. The remaining high-quality alignments were assigned to genes using featureCounts function from Bioconductor Rsubread package version 2.12.3 with inbuilt mm39 annotation. Differential Expression along with adjusted p-values (FDR) and log Fold Changes (logFC) were computed using Bioconductor edgeR package (glmLRT function). Immunoglobulin genes are highly expressed in plasma cells, and thus were excluded from the analysis. Genes were deemed differentially expressed if their FDR was less than 0.05 and Fold Change greater than 2 (|logFC| > 1). *Adam6a* is the most highly upregulated gene in ASCs following PTC-028 treatment, however, it is positioned in the V-D region and has the opposite orientation to the immunoglobulin genes (45). It is likely this sequence is being transcribed because of immunoglobulin gene transcription and has been misaligned during processing and thus was removed from further analysis. Volcano plots were made using ggVolcanoR (46).

#### GO analysis

Selected genes were included in GO analysis if they had an FDR < 0.10 and Fold Change > 1.5 (|logFC| > log2(1.5)). We further split into genes upregulated and downregulated in PTC-028 treated vs Control samples, and GO analysis performed for each subset separately. Bioconductor topGO package version 2.50.0 with classic algorithm and Exact Fisher Test was used.

#### Cell Cycle Enrichment Analysis

We used the list of cell cycle related genes from (47) and converted them into mouse genes where applicable. We used the Enrichment Score method as in (48) separately taking G2/M genes, G1/S and all of the above as our Gene Set. Since there were 4 samples, we could not permute the sample labels, so we permuted the genes. Based on 1 billion permutations, the maximum deviations were 0.57, 0.54 and 0.50 respectively, lower than the ES, so based on these tests the p-values are < 10^-9^ for each set.

### Human ASC survival culture

ASCs (FVS780^-^CD20^-^CD3^-^CD14^-^CD16^-^CD19^+^CD27^hi^CD38^hi^) were sort-purified from human peripheral blood mononuclear cells into culture media (10% FCS, 2 mM L-glutamine (Gibco), 100 units/mL Penicillin and 100 ng/mL Streptomycin (Sigma-Aldrich), 10 mM HEPES (Gibco), 1 mM Sodium Pyruvate (Gibco) in RPMI). Two of the biological replicates within each group (patients or healthy donors) were each obtained from pooling sort-purified ASCs from two individual donors, due to low cell number. Samples were pooled following sort-purification. The primary human tonsil fibroblast line was a kind gift from Shannon Turley and has been previously characterised (49). It was used at passage 4-X. Fibroblasts were cultured in the above media for 3-4 days in a 75 cm^3^ sterile culture flask (Falcon) or until confluent. During passaging of human tonsil fibroblasts, supernatant was kept and frozen. Fibroblast supernatant batches that promoted high survival of ASCs were pooled and used for the survival assay. 3000 – 5000 ASCs per well were cultured in sterile 96-well tissue culture plates (Falcon) in the presence of 20 ng/mL IL-6 (R&D Systems), 100 ng/mL APRIL (R&D Systems) diluted in supernatant from human tonsillar fibroblasts. ASCs were cultured for 6 days in the presence of 1.25 μM PTC-028 dissolved in DMSO (Sigma) or DMSO only as a vehicle control. 10^4^ Calibrite Beads (BD) were added per well to facilitate enumeration of cells. Cells were harvested and viability and surface marker expression determined via flow cytometry. The number of live ASCs was calculated based on the known number of beads added per well and normalised for the number of cells initially cultured. Collection and/or use of human cells was ethically reviewed and approved by the Monash University Human Research Ethics Committee.

### Statistical analysis

Mann-Whitney nonparametric two tailed tests and ratio paired t tests were used for statistical analysis. * p≤0.05, ** p≤0.01, or exact p values are stated with p≤0.05 treated as significant. Data are presented as mean +/- standard error of the mean (SEM) and/or individual data points. Analysis was performed using GraphPad Prism v9 (GraphPad Software).

## Acknowledgments

We thank Zoe Chua for technical assistance, Liam Kealy for critical reading of this manuscript, and staff of the Monash University FlowCore, Animal Research Platform.

---

**Supplementary Figure 1:**
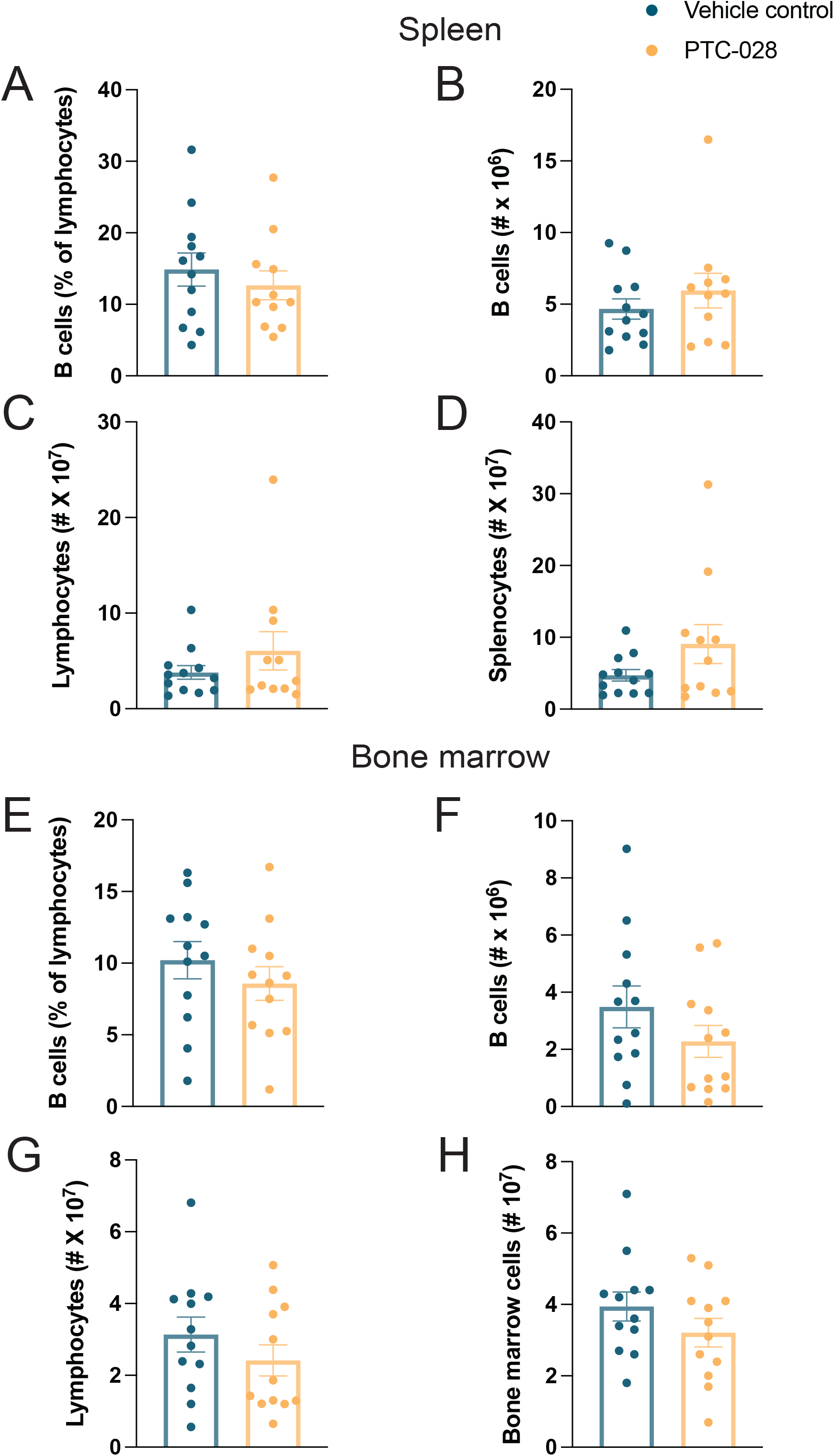
PTC-028 did not alter B cell, lymphocytes or total cellularity in the spleen on bone marrow. *Lyn^-/-^* mice were treated with vehicle control or PTC-028 and assessed on d15. A) Frequency of splenic B cells as a proportion of total lymphocytes; B) total number of splenic B cells; C) total number of lymphocytes in the spleen; D) total number of splenocytes. E) Frequency of bone marrow B cells as a proportion of total lymphocytes; F) total number of bone marrow B cell; G) total number of lymphocytes in the bone marrow; D) total number of bone marrow cells. Data is presented as the mean +/- SEM with individual data points representing one mouse. Data was combined from three independent experiments. n = 11–12 per group. Mann-Whitney nonparametric two tailed tests were used for statistical analysis.

**Supplementary Figure 2:**
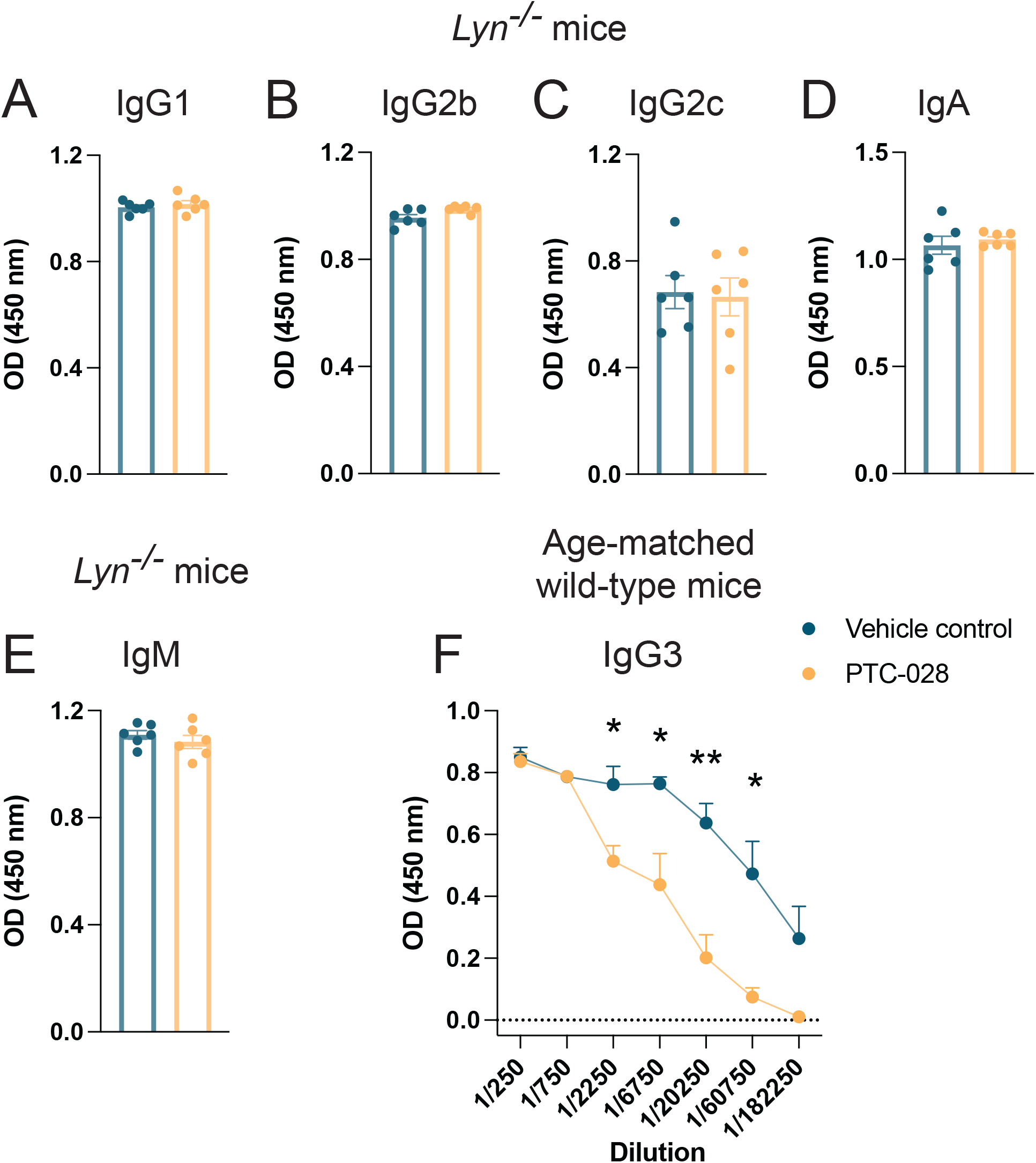
Assessment of serum antibody in *Lyn^-/-^* mice and age-matched wild type mice. A) IgG1, B) IgG2b, C) IgG2c, D) IgA and E) IgM in *Lyn^-/-^*mice treated with vehicle control or PTC-028 once daily on d0 - 11 and then assessed on d15. Data shown is representative of three independent experiments. n=6 per group. Data is presented as the mean +/- SEM with individual data points representing one mouse. F) ELISA results measuring total serum IgG3 antibody from wild-type mice (≥ 6 months old) on d15. Data shown is representative of two independent experiments. n=5 per group. Data is presented as the mean +/- SEM. Mann-Whitney nonparametric two tailed tests was used for statistical analysis. *P≤0.05, **P≤0.01.

**Supplementary Table 1:** Sjögren’s Syndrome patient details.

| Code ID | Sex | Age | Year of diagnosis | Sjogren's manifestations | Comorbidities | Anti-SSA | RF | Serum IgG (g/L, NR 6-16) | ESSDAI score | Medication |
| --- | --- | --- | --- | --- | --- | --- | --- | --- | --- | --- |
| SS01 | F | 32 | 2018 | Sicca<br>Salivary gland enlargement<br>Annular erythema<br>Lymphopenia | Pituitary microadenoma<br>Autoimmune hypothyroidism | Yes | Yes | 16.75 | 3 | Prednisolone 3mg<br>Hydroxychloroquine |
| SS02 | F | 60 | 2019 | Sicca<br>Inflammatory arthritis | Colonic adenocarcinoma<br>Malignant melanoma | Yes | Yes | 14.29 | 2 | Meloxicam<br>Oroxine<br>Propranolol<br>Relpax<br>Rosuvastatin<br>Topiramate |
| SS03 | F | 61 | 2019 | Sicca<br>Salivary gland enlargement | Thyroid carcinoma<br>Parotid gland mucoepidermoid carcinoma<br>Type 2 diabetes mellitus<br>Diabetic peripheral neuropathy | Yes | Not done | 10.78 | 2 | Januvia<br>Metformin<br>Thyroxine<br>Rosuvastatin<br>Reni |
| SS04 | F | 62 | 2017 | Sicca<br>Lymphocytic interstitial pneumonitis | Obstructive sleep apnoea<br>Hypertension<br>Autoimmune hypothyroidism | Yes | Yes | 24.04 | 26 | Hydroxychloroquine<br>Thyroxine<br>Olmesartan<br>Diclofenac<br>Nexium |
|  |  |  |  | Cutaneous lupus<br>erythematosus<br>Inflammatory<br>arthritis<br>Lymphopenia |  |  |  |  |  | Paracetamol<br>Nocturnal oxygen |
| SS05 | F | 62 | 2017 | Sicca<br>Peripheral and<br>autonomic<br>neuropathy<br>Lymphopenia | Colonic carcinoma<br>Hypertension<br>Type 2 diabetes<br>mellitus<br>Exocrine pancreatic<br>insufficiency | Yes | Yes | 8.4 | 13 | IVIg<br>Hydroxychloroquine<br>Amphotericin<br>Amgapre<br>Ondansetron<br>Tramadol<br>Paracetamol<br>Atorvastatin<br>Esomeprazole<br>Lercadip<br>Metformin<br>Pritor Plus<br>Carbamgapre |
| SS06 | F | 67 |  | Sicca<br>Salivary gland<br>enlargement<br>Lymphadenopat-<br>hy<br>Splenomegaly<br>Cutaneous<br>vasculitis<br>Lymphocytic<br>interstitial<br>pneumonitis | Lymphocytic<br>interstitial<br>pneumonia | Yes | Yes | 10.8 | 25 | Prednisolone 10mg<br>Mycophenolate<br>mofetil<br>Rituximab |
|  |  |  |  | Inflammatory<br>arthritis<br>Anaemia |  |  |  |  |  |  |

**Supplementary Table 2:** Age and sex matched healthy donor details.

| Code ID | Sex | Age | Patient match |
| --- | --- | --- | --- |
| HD01 | Female | 33 | SS01 |
| HD02 | Female | 61 | SS02 |
| HD03 | Female | 60 | SS03 |
| HD04 | Female | 64 | SS04 |
| HD05 | Female | 63 | SS05 |
| HD06 | Female | 67 | SS06 |

## REFERENCES

1. Barber MRW, Drenkard C, Falasinnu T, et al. Global epidemiology of systemic lupus erythematosus. Nature Reviews Rheumatology. 2021;17(9):515–32.

2. Lech M, Anders H-J. The Pathogenesis of Lupus Nephritis. Journal of the American Society of Nephrology. 2013;24(9):1357–66.

3. Patel R, Shahane A. The epidemiology of Sjogren’s syndrome. Clin Epidemiol. 2014;6:247–55.

4. Mielle J, Tison A, Cornec D, Le Pottier L, Daien C, Pers JO. B cells in Sjogren’s syndrome: from pathophysiology to therapeutic target. Rheumatology (Oxford). 2021;60(6):2545–60.

5. Verstappen GM, Pringle S, Bootsma H, Kroese FGM. Epithelial-immune cell interplay in primary Sjogren syndrome salivary gland pathogenesis. Nat Rev Rheumatol. 2021;17(6):333–48.

6. Huang H, Benoist C, Mathis D. Rituximab specifically depletes short-lived autoreactive plasma cells in a mouse model of inflammatory arthritis. Proceedings of the National Academy of Sciences. 2010;107(10):4658–63.

7. Marcondes F, Scheinberg M. Belimumab in the treatment of systemic lupus erythematous: An evidence based review of its place in therapy. Autoimmunity reviews. 2018;17(2):103–7.

8. Mumtaz IM, Hoyer BF, Panne D, et al. Bone marrow of NZB/W mice is the major site for plasma cells resistant to dexamethasone and cyclophosphamide: Implications for the treatment of autoimmunity. Journal of Autoimmunity. 2012;39(3):180–8.

9. Vallerskog T, Gunnarsson I, Widhe M, et al. Treatment with rituximab affects both the cellular and the humoral arm of the immune system in patients with SLE. Clin Immunol. 2007;122(1):62–74.

10. Alexander T, Cheng Q, Klotsche J, et al. Proteasome inhibition with bortezomib induces a therapeutically relevant depletion of plasma cells in SLE but does not target their precursors. European Journal of Immunology. 2018;48(9):1573–9.

11. Kealy L, Di Pietro A, Hailes L, et al. The histone methyltransferase DOT1L is essential for humoral immune responses. Cell reports. 2020;33(11):108504.

12. Béguelin W, Popovic R, Teater M, et al. EZH2 Is Required for Germinal Center Formation and Somatic EZH2 Mutations Promote Lymphoid Transformation. Cancer cell. 2013;23(5):677–92.

13. Di Pietro A, Polmear J, Cooper L, et al. Targeting BMI-1 in B cells restores effective humoral immune responses and controls chronic viral infection. Nat Immunol. 2022;23(1):86–98.

14. Caganova M, Carrisi C, Varano G, et al. Germinal center dysregulation by histone methyltransferase EZH2 promotes lymphomagenesis. J Clin Invest. 2013;123(12):5009–22.

15. Good-Jacobson KL, Chen Y, Voss AK, Smyth GK, Thomas T, Tarlinton D. Regulation of germinal center responses and B-cell memory by the chromatin modifier MOZ. Proc Natl Acad Sci U S A. 2014;111(26):9585–90.

16. Imhof A. Epigenetic regulators and histone modification. Briefings in Functional Genomics. 2006;5(3):222–7.

17. Zauderer MG, Szlosarek PW, Le Moulec S, et al. EZH2 inhibitor tazemetostat in patients with relapsed or refractory, BAP1-inactivated malignant pleural mesothelioma: a multicentre, open-label, phase 2 study. Lancet Oncol. 2022;23(6):758–67.

18. Han M, Jia L, Lv W, Wang L, Cui W. Epigenetic Enzyme Mutations: Role in Tumorigenesis and Molecular Inhibitors. Frontiers in Oncology. 2019;9(194).

19. Di Pietro A, Good-Jacobson KL. Disrupting the Code: Epigenetic Dysregulation of Lymphocyte Function during Infectious Disease and Lymphoma Development. J Immunol. 2018;201(4):1109–18.

20. Guo M, Price MJ, Patterson DG, et al. EZH2 Represses the B Cell Transcriptional Program and Regulates Antibody-Secreting Cell Metabolism and Antibody Production. J Immunol. 2018;200(3):1039–52.

21. Tarte K, Zhan F, De Vos J, Klein B, Shaughnessy J, Jr. Gene expression profiling of plasma cells and plasmablasts: toward a better understanding of the late stages of B-cell differentiation. Blood. 2003;102(2):592–600.

22. Hibbs ML, Tarlinton DM, Armes J, et al. Multiple defects in the immune system of <em>Lyn</em>-deficient mice, culminating in autoimmune disease. Cell. 1995;83(2):301–11.

23. Silver KL, Crockford TL, Bouriez-Jones T, Milling S, Lambe T, Cornall RJ. MyD88-dependent autoimmune disease in Lyn-deficient mice. Eur J Immunol. 2007;37(10):2734–43.

24. Tsantikos E, Oracki SA, Quilici C, Anderson GP, Tarlinton DM, Hibbs ML. Autoimmune disease in Lyn-deficient mice is dependent on an inflammatory environment established by IL-6. J Immunol. 2010;184(3):1348–60.

25. Dey A, Xiong X, Crim A, et al. Evaluating the Mechanism and Therapeutic Potential of PTC-028, a Novel Inhibitor of BMI-1 Function in Ovarian Cancer. Mol Cancer Ther. 2018;17(1):39–49.

26. DiLillo DJ, Hamaguchi Y, Ueda Y, et al. Maintenance of long-lived plasma cells and serological memory despite mature and memory B cell depletion during CD20 immunotherapy in mice. J Immunol. 2008;180(1):361–71.

27. Lau M, Tsantikos E, Maxwell MJ, Tarlinton DM, Anderson GP, Hibbs ML. Loss of STAT6 promotes autoimmune disease and atopy on a susceptible genetic background. J Autoimmun. 2012;39(4):388–97.

28. Park IK, Morrison SJ, Clarke MF. Bmi1, stem cells, and senescence regulation. J Clin Invest. 2004;113(2):175–9.

29. Jacobs JJ, Kieboom K, Marino S, DePinho RA, van Lohuizen M. The oncogene and Polycomb-group gene bmi-1 regulates cell proliferation and senescence through the ink4a locus. Nature. 1999;397(6715):164–8.

30. Behesti H, Bhagat H, Dubuc AM, Taylor MD, Marino S. Bmi1 overexpression in the cerebellar granule cell lineage of mice affects cell proliferation and survival without initiating medulloblastoma formation. Dis Model Mech. 2013;6(1):49–63.

31. Lin X, Wei F, Whyte P, Tang D. BMI1 reduces ATR activation and signalling caused by hydroxyurea. Oncotarget. 2017;8(52):89707–21.

32. Obuse C, Yang H, Nozaki N, Goto S, Okazaki T, Yoda K. Proteomics analysis of the centromere complex from HeLa interphase cells: UV-damaged DNA binding protein 1 (DDB-1) is a component of the CEN-complex, while BMI-1 is transiently co-localized with the centromeric region in interphase. Genes Cells. 2004;9(2):105–20.

33. Szyszko EA, Brun JG, Skarstein K, Peck AB, Jonsson R, Brokstad KA. Phenotypic diversity of peripheral blood plasma cells in primary Sjogren’s syndrome. Scand J Immunol. 2011;73(1):18–28.

34. Steinmetz TD, Verstappen GM, Schulz SR, et al. Association of Circulating Antibody-Secreting Cell Maturity With Disease Features in Primary Sjogren’s Syndrome. Arthritis Rheumatol. 2023;75(6):973–83.

35. Antonis F, Myrto K, Alessia A, et al. 2019 update of the EULAR recommendations for the management of systemic lupus erythematosus. Annals of the Rheumatic Diseases. 2019;78(6):736.

36. Tanaka Y, Nakayamada S, Yamaoka K, Ohmura K, Yasuda S. Rituximab in the real-world treatment of lupus nephritis: A retrospective cohort study in Japan. Modern Rheumatology. 2022;33(1):145–53.

37. Manfre V, Cafaro G, Riccucci I, et al. One year in review 2020: comorbidities, diagnosis and treatment of primary Sjogren’s syndrome. Clin Exp Rheumatol. 2020;38 Suppl 126(4):10–22.

38. Grigoriadou S, Chowdhury F, Pontarini E, Tappuni A, Bowman SJ, Bombardieri M. B cell depletion with rituximab in the treatment of primary Sjogren’s syndrome: what have we learnt? Clin Exp Rheumatol. 2019;37 Suppl 118(3):217–24.

39. Cao R, Tsukada Y-i, Zhang Y. Role of Bmi-1 and Ring1A in H2A Ubiquitylation and Hox Gene Silencing. Molecular Cell. 2005;20(6):845–54.

40. Klein-Schneegans AS, Kuntz L, Fonteneau P, Loor F. Serum concentrations of IgM, IgG1, IgG2b, IgG3 and IgA in C57BL/6 mice and their congenics at the lpr (lymphoproliferation) locus. J Autoimmun. 1989;2(6):869-75.

41. Klaus T, Bereta J. CH2 Domain of Mouse IgG3 Governs Antibody Oligomerization, Increases Functional Affinity to Multivalent Antigens and Enhances Hemagglutination. Front Immunol. 2018;9:1096.

42. Bruhns P. Properties of mouse and human IgG receptors and their contribution to disease models. Blood. 2012;119(24):5640–9.

43. Klaus T, Bzowska M, Kulesza M, et al. Agglutinating mouse IgG3 compares favourably with IgMs in typing of the blood group B antigen: Functionality and stability studies. Sci Rep. 2016;6:30938.

44. Greenspan NS, Lu MA, Shipley JW, et al. IgG3 deficiency extends lifespan and attenuates progression of glomerulonephritis in MRL/lpr mice. Biol Direct. 2012;7:3.

45. Featherstone K, Wood AL, Bowen AJ, Corcoran AE. The mouse immunoglobulin heavy chain V-D intergenic sequence contains insulators that may regulate ordered V(D)J recombination. J Biol Chem. 2010;285(13):9327–38.

46. Mullan KA, Bramberger LM, Munday PR, et al. ggVolcanoR: A Shiny app for customizable visualization of differential expression datasets. Comput Struct Biotechnol J. 2021;19:5735–40.

47. Subramanian A, Tamayo P, Mootha VK, et al. Gene set enrichment analysis: a knowledge-based approach for interpreting genome-wide expression profiles. Proc Natl Acad Sci U S A. 2005;102(43):15545–50.

48. Fischer M, Grossmann P, Padi M, DeCaprio JA. Integration of TP53, DREAM, MMB-FOXM1 and RB-E2F target gene analyses identifies cell cycle gene regulatory networks. Nucleic Acids Res. 2016;44(13):6070–86.

49. Knoblich K, Cruz Migoni S, Siew SM, et al. The human lymph node microenvironment unilaterally regulates T-cell activation and differentiation. PLoS Biol. 2018;16(9):e2005046.

